# A novel essential RNA-binding protein complex in *Trypanosoma brucei*

**DOI:** 10.1101/2020.05.19.104984

**Authors:** Kathrin Bajak, Kevin Leiss, Christine Clayton, Esteban Erben

## Abstract

ZC3H5 is an essential cytoplasmic trypanosome protein with a single Cx_7_Cx_5_Cx_3_H zinc finger domain. We here show that ZC3H5 forms a complex with three other proteins, encoded by genes Tb927.11.4900, Tb927.8.1500 and Tb927.7.3040. ZC3H5 interacts directly with Tb927.11.4900, which in turn interacts with Tb927.7.3040. Tb927.11.4900 has a circularly permuted GTPase domain, which is required for the Tb927.7.3040 interaction. RNA immunoprecipitation revealed that ZC3H5 is preferentially associated with poorly translated, low-stability mRNAs, the 5’-untranslated regions and coding regions of which are enriched in the motif (U/A)UAG(U/A). Tethering of ZC3H5, or other complex components, to a reporter repressed its expression. However, depletion of ZC3H5 *in vivo* did not increase the abundance of ZC3H5-bound mRNAs: instead, counter-intuitively, there were very minor decreases in a few targets, and marked increases in the abundances of very stable mRNAs encoding ribosomal proteins. Depletion also resulted in an increase in monosomes at the expense of large polysomes, and appearance of “halfmer” disomes containing two 80S subunits and one 40S subunit. We speculate that the ZC3H5 complex might be implicated in quality control during the translation of sub-optimal open reading frames; complex assembly might be regulated by GTP hydrolysis and GTP-GDP exchange.

---

*Trypanosoma brucei* is a unicellular eukaryote that proliferates in the blood and tissue fluids of mammals and in the digestive system of Tsetse flies. *T. brucei* and related trypanosomes cause African sleeping sickness in humans and nagana in cattle: diseases which produce a significant economic burden for a vast region of Africa. Two *T. brucei* life cycle stages are easily cultured in the laboratory: bloodstream forms (BF) can be grown in high-glucose medium at 37°C whereas procyclic forms (PF) are cultivated in high-proline medium at 27°C.

Kinetoplastids rely almost exclusively on post-transcriptional mechanisms for control of gene expression. Transcription is polycistronic and individual mRNAs are generated by trans splicing of a 39nt leader to the 5’-end, and by 3’ polyadenylation (1). Trypanosome mRNAs vary extensively in decay rates and translation efficiency. Work by numerous laboratories has demonstrated that RNA-binding proteins (RBPs) play prominent roles in the regulation of splicing, translation and mRNA decay, although the mechanisms by which they do this have - with a few exceptions - remained obscure (1).

Two high-throughput approaches enabled us to identify RBPs that can increase or decrease expression of bound mRNAs. The tethering assay involves co-expression of the protein of interest fused to a highly sequence-specific RNA-binding peptide, and a reporter mRNA bearing the cognate recognition sequence; we use the lambdaN peptide - boxB combination. We conducted genome-wide library screens to reveal protein fragments that can either activate or repress expression of a reporter when tethered (2); results that were later verified for a subset of full-length proteins (3). In addition, a catalogue of proteins that can be cross-linked to BF mRNAs (the trypanosome “mRNP proteome”) was obtained (3). This approach led to the identification of 155 high-confidence RBPs carrying both canonical (RRM, PUF and Zinc finger) and potential novel RNA-binding domains. Although several of the identified RBPs are essential for stage-specific gene expression and/or have developmentally regulated expression or modification (reviewed in (9)), only a small fraction of them exhibited a clear effect on expression when tethered to the mRNA reporter.

One RBP that did have an effect when tethered is a cytoplasmic zinc finger protein named ZC3H5 (Tb927.3.740) (2,3). ZC3H5 can be cross-linked to, and purifies with, poly(A)+ RNA (3), it decreases gene expression when tethered to a reporter (3), its expression is similar in multiplying BF and PF trypanosomes, and high-throughput RNAi analysis suggested that it is essential in BF (4). In this study, we investigated the role of ZC3H5 in regulating gene expression in *T. brucei.* We found that ZC3H5 is part of a protein complex which is potentially regulated by GTP binding. RNAi-mediated depletion of ZC3H5 or any other member of the complex results in cytokinesis inhibition. While ZC3H5 preferentially associates with poorly translated, low-stability mRNAs, its depletion results in only minor changes to the transcriptome. We hypothesize that ZC3H5 is involved in quality control during the translation of sub-optimal transcripts.

## Results

### Conservation of ZC3H5 in Kinetoplastea

ZC3H5 is a 25.5 kDa protein with a single Cx7Cx5Cx3H zinc finger domain (Fig. 1A). ZC3H5 is present throughout the Kinetoplastea but absent in Euglenids (Supplementary Fig. S1). Comparison of sequences from *Bodo, Paratrypanosoma, Crithidia, Endotrypanum, Leishmania* and *Trypanosoma* reveals that the 52 residues surrounding the CCCH domain (5 residues N-terminal, 21 C-terminal) are 51% identical and 81% similar; conservation is stronger if *Bodo* and *Paratrypanosoma* are excluded. The N-termini lack charged residues, while the C-termini are relatively proline-rich. *Paratrypanosoma* ZC3H5 has an 800-residue C-terminal extension.

**Fig 1.**
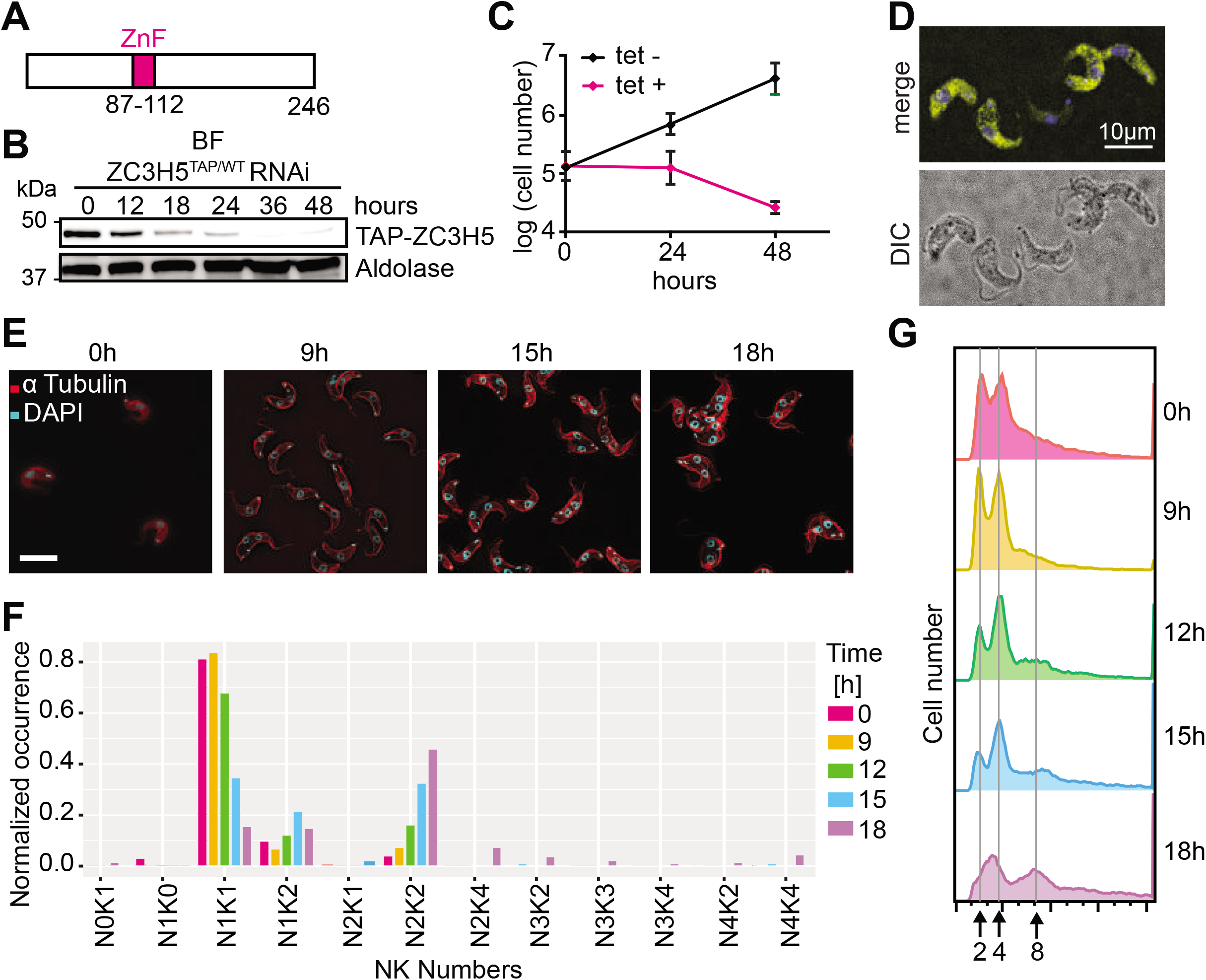
Depletion of ZC3H5 inhibits cytokinesis. (A) Structure of ZC3H5, to scale. The CCCH zinc finger, as detected by SMART, is shown in magenta. (B) Disappearance of TAP-ZC3H5 after RNAi. In the cell line used, one copy was in situ TAP-tagged and the other was intact. Aldolase served as a control. The numbers of hours after tetracycline addition to induce RNAi are shown. (C) Effect of RNAi on cell numbers. Cumulative cell numbers with and without tetracycline addition are shown for 3 replicates, with standard deviation. (D) YFP-ZC3H5 is in the cytoplasm. Bloodstream-form trypanosomes expressing YFP-ZC3H5 from the endogenous locus were stained for YFP (yellow) and for DNA with DAPI (blue). Images were examined using the Leica DMi8 spinning disk microscope (Scale bar: 10 μm). (E) Effect of ZC3H5 depletion on cell morphology. Cells were stained for DNA (blue) and tubulin (red). (F) The number of nuclei (N) and kinetoplasts (K) per cell of the population described in E was quantified (n>200) at the time-points indicated. (G) Effect of ZC3H5 depletion on DNA content. DNA was stained with Propidium iodide and the cells were analyzed by FACS. The numbers below the diagrams indicate ploidy.

### Depletion of ZC3H5 inhibits cytokinesis

We first confirmed that ZC3H5 is indeed required for BF trypanosome proliferation. To enable detection, we integrated a tandem affinity purification tag sequence (protein A - calmodulin binding peptide) upstream of, and in frame with, one ZC3H5 open reading frame (TAP-ZC3H5). Induction of RNA interference caused a reduction in TAP-ZC3H5 expression within 12h (Fig 1B), and inhibited cell proliferation (Fig 1C). Morphological analysis (Fig 1E, F) and DNA content measurements (Fig 1G) revealed that the cells were incapable of cytokinesis. Within 18h of RNAi induction, multinucleate cells were beginning to appear. The DNA profile of cells without RNAi induction suggested the presence of more 4N cells than normal; this perhaps suggests that the cells have somewhat less ZC3H5 than normal. We also verified the cytoplasmic location of ZC3H5, using a cell line with N-terminally *in situ* YFP-tagged protein (Fig 1D).

### ZC3H5 recruits a protein complex

A two-hybrid screen using CAF1 deadenylase as bait suggested an interaction with ZC3H5 (3). To examine this and other possible interactions, we purified TAP-tagged ZC3H5 and subjected the preparations to mass spectrometry (MS). In order to find proteins that were specific to ZC3H5, we compared the results with those from a similar purification of ERBP1 (5). (Essentially same results were obtained TAP-GFP). The results revealed robust co-purification of ZC3H5 with the proteins encoded by Tb927.11.4900, Tb927.7.3040, and Tb927.8.1500 (Fig 2A; Supplementary Table S1). Co-precipitations of the three newly-found proteins combined with MS analysis confirmed that they indeed form a complex (Fig 2B-D). ZC3H5 was however not significantly enriched, suggesting that only a small proportion of 11.4900-7.3040-8.1500 complexes is interacting with ZC3H5. Nevertheless, pull-down of V5-*in-situ*-tagged versions of Tb927.11.4900, Tb927.8.1500 and Tb927.7.3040 resulted in clear coimmunoprecipitation of some tagged ZC3H5 (Fig 2E, F), although most remained in the unbound fraction. The results therefore indicated that a subpopulation of ZC3H5 molecules recruits the Tb927.11.4900-Tb927.8.1500-Tb927.7.3040 complex. All three proteins were in the bloodstream-form mRNP proteome (3) (FDR= 0.026, 0.0018 and 0.011, respectively) and are cytoplasmic when fused to GFP (6).

**Fig 2.**
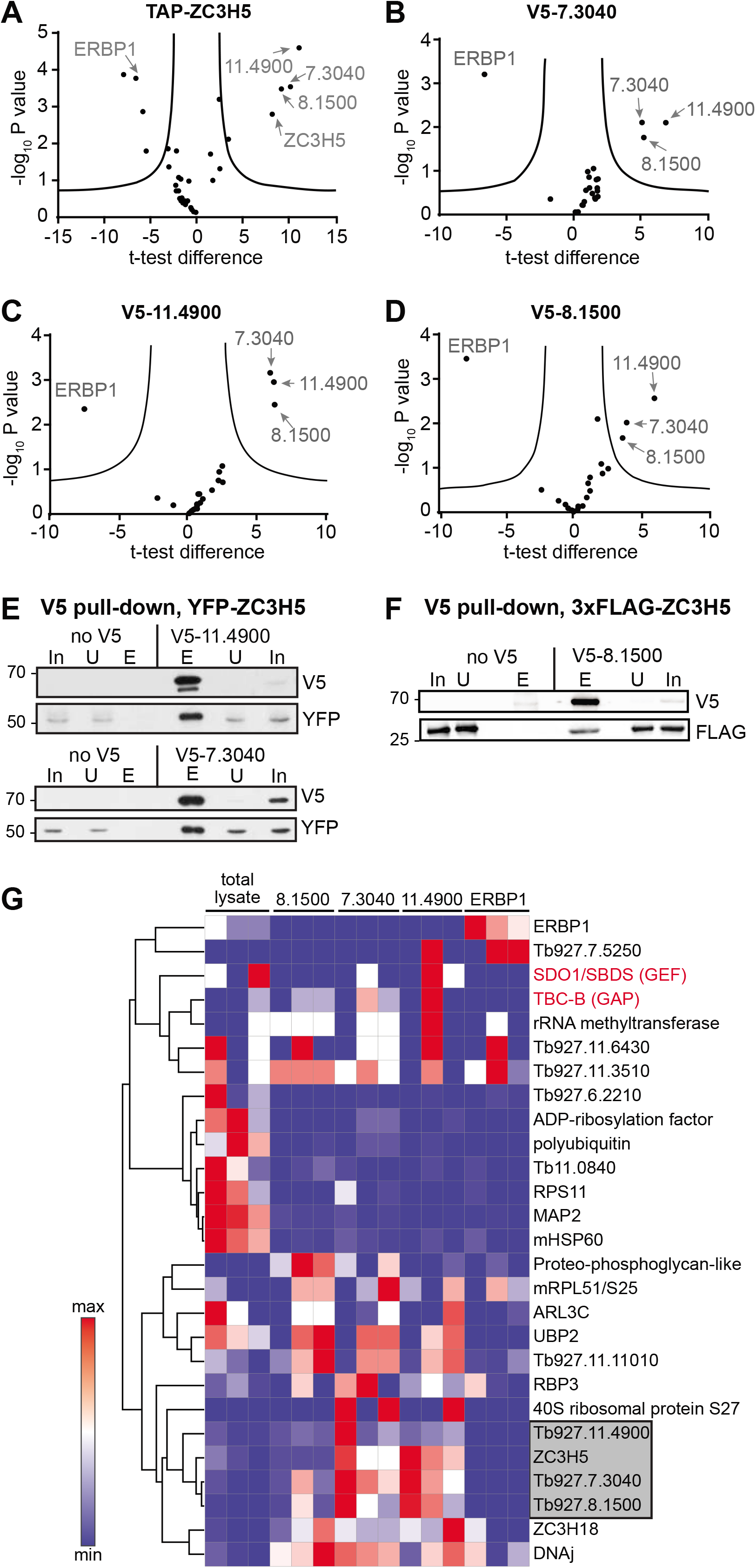
ZC3H5 is associated with three other proteins. (A) Endogenously TAP-tagged ZC3H5 was purified three times. Raw data were analyzed by MaxQuant, and specific interactors were selected from background using label-free quantification in Perseus. In the volcano plots, the ratio of the results with TAP-ZC3H5 to a different TAP-tagged protein, ERBP1, are shown on the x-axis, and the log10 of the false discovery rate (FDR), calculated by a permutation-based FDR adapted t-test, is on the y-axis. Significantly selected proteins are labeled. Gene numbers are truncated by removal of “Tb927.” Data are in Supplementary Table S1. (B) As (A) but for V5-Tb927.7.3040. (C) As (A) but for V5-Tb927.11.4900. (D) As (A) but for V5-Tb927.8.1500. (E) The cells used expressed YFP-ZC3H5 and V5-tagged versions of the interaction partners Tb927.7.3040 and Tb927.11.4900. After immunoprecipitation the proteins were detected by Western blotting. The eluates are from 10 times more cells than were used for the input and unbound lanes. (F) As (E), except that the cells used expressed 3xFLAG-ZC3H5 and V5-tagged Tb927.8.1500. (G) Additional interaction partners of the ZC3H5 complex. The heat map compares triplicate results after purification of V5-tagged versions of Tb927.8.1500, Tb927.7.3040, and Tb927.11.4900, or ERBP1, and includes only proteins that were detected in at least two replicates. Results for total lysate are also included as an additional control. ZC3H5 complex components are shaded in grey. The trypanosome homologue of *Saccharomyces cerevisiae* Sdo1 (human SBDS), a likely guanine nucleotide exchange factor (Tb927.4.3140), and TBC-B (Tb927.7.2470), a likely GTPase activating factor, are highlighted in red.

Interestingly, scrutiny of the mass spectrometry results (Fig 2G) revealed weak association with a cytoplasmic protein that is probably a homologue of yeast GTP-GDP exchange factor Sdo1 (Tb927.4.3140, BLAST E-value 4e-45). There was also a putative cytoplasmic GTPase activating protein, Tb927.7.2470. We speculate that these two proteins might be involved in regulating formation of the ZC3H5 complex.

### Characteristics of the Tb927.7.3040, Tb927.8.1500 and Tb927.11.4900proteins

We examined the characteristics of the newly-identified proteins by comparing the sequences in selected representatives of Trypanosomatida, as well as the more distantly-related Kinetoplastid (class Kinetoplastea) *Bodo saltans* (7). Tb927.7.3040 is a 69 kDa protein which has C-terminal WD repeats. It was localized to starvation stress granules in procyclic forms (8) and is conserved as far as *Bodo saltans* (Supplementary Fig S2). The N-terminus belongs to the PTHR19924 family, which contains various proteins implicated in rRNA processing.

The 63 kDa Tb927.8.1500 protein is conserved throughout Kinetoplastea (Supplementary Fig S3). The sequence alignment reveals three different ~30 residue conserved regions starting at the extreme N-terminus and at residues 170 and 262 (*T. brucei* sequence numbers) (Supplementary Fig S3). The C-terminal regions are very variable and include proline- and glutamine-rich tracts.

Tb927.11.4900 is absent in *Bodo* but present in Trypanosomatida. It is a 62 kDa protein and contains four WD40 repeats towards the C-terminus (Supplementary Fig S4). Pro-site scans classify it as being a member of the “F-box and WD40 family” (PTHR44156), but it lacks an F-box. PTHR44156 includes some guanine nucleotide binding beta subunits, and indeed the region of Tb927.11.4900 upstream of the WD repeats includes (in this order) G4, G1 and G3 P-loop NTPase motifs (Fig 3A, Supplementary Fig S4). This order is similar to that seen in eukaryotic and prokaryotic circularly permutated GTPases, which have the motif order G4-G5’-G1-G2’-G3 (9,10). The G4 domain (N/T)KxD, which determines specificity for guanine, is present in *T. brucei* Tb927.11.4900 and changed to TKxE in other trypanosomes (which should not prevent activity). However, it is less well conserved in other kinetoplastids (Supplementary Fig S4). The G1 (P-loop) motif in Tb927.11.4900 has leucine instead of serine/threonine but the lysine required for phosphate interaction is conserved; while the G3 domain (Walker B motif), (D/E)xxG, which binds Mg^2+^ and is required for tight nucleotide binding and hydrolysis, is conserved throughout. The G5’ and G2’ motifs are generally poorly conserved and difficult to recognize in primary sequences; G2 has only one threonine totally conserved (10). Threonines are present between the putative G1 and G3 domains in all species examined except *Paratrypanosoma* (Supplementary Fig S4).

**Fig 3.**
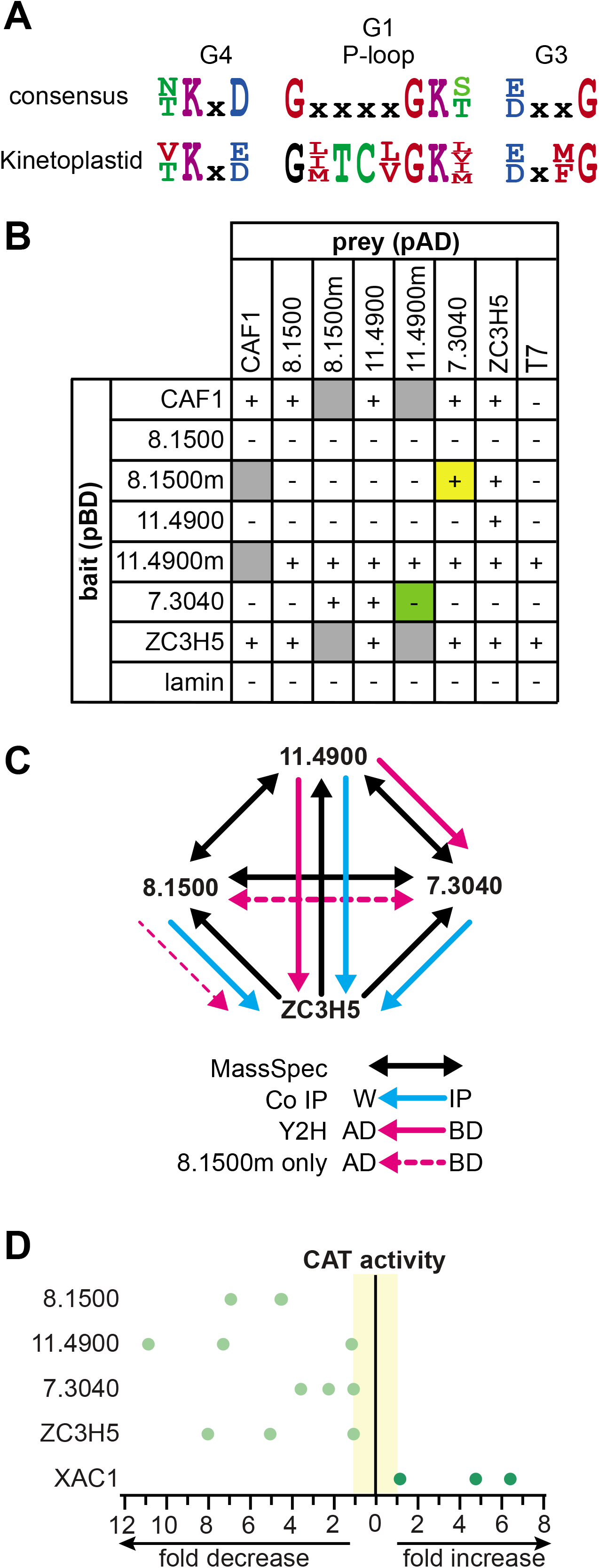
Characteristics and interactions of ZC3H5 complex proteins. (A) The consensus G4, G1 and G3 sequence motifs as described in (10) are shown, with the equivalents from Tb927.11.4900 and its homologues. (B) Results from yeast 2-hybrid analysis. “+” indicates that the interactions were positive both by growth and alpha-galactosidase expression. Interactions that were not tested are shown in grey. Note that ZC3H5 and Tb927.11.4900m were auto-activators when fused to the DNA-binding domain. Tb927.8.1500m has a T443E and Tb927.11.4900m has TKVD/AKVA. (C) Summary of interaction studies. The results suggest that Tb927.11.4900 is a central component that interacts with both ZC3H5 and Tb927.7.3040. Tb927.8.1500 interacts with Tb927.7.3040 only when a phospho-mimic mutation (T443E) is present in Tb927.8.1500. (D) All four proteins decrease expression when tethered. Expression of lambdaN-test-protein-myc was induced for 24h then CAT activity was measured relative to the activity in the absence of induction. Tb927.7.2780 (XAC1) served as a positive control.

### Interactions between the complex components

To examine interactions between the complex components we used pair-wise yeast two-hybrid assays. ZC3H5 was a constitutive activator as bait, and activity of CAF1 bait was also promiscuous (Fig 3B, Supplementary Fig S5). The results for ZC3H5 as prey, however, suggested that it interacts directly with Tb927.11.4900. There was also an interaction between Tb927.7.3040 (bait) and Tb927.11.4900 (prey) although the reciprocal interaction was not observed. We concluded that ZC3H5 recruits Tb927.7.3040 via Tb927.11.4900 (Fig 3C). No direct interaction of Tb927.8.1500 was observed. However, Tb927.8.1500 has four phosphorylated serines (11,12) and three phosphothreonines (Supplementary Fig S3), one of which, T443, shows peak phosphorylation in late S phase (12). To investigate the effect of the regulated phosphorylation we mutated the relevant threonine (arrow in Supplementary Fig S3) to aspartate as a phospho-mimic. Interestingly, reciprocal interactions were observed between Tb927.8.1500 T443D and Tb927.7.3040, suggesting that full complex formation might be cell-cycle regulated (Fig 3B, yellow highlight). However, preliminary coimmunoprecipitation results (not shown) indicated that association of Tb927.8.1500 with ZC3H5 was present in both T443D and T443A mutants.

To investigate the role of the putative GTP-binding domain in Tb927.11.4900 we mutated the G4 motif, TKVD, to AKVA. In the yeast two-hybrid assay, the resulting protein became constitutively active as bait, but as prey, the interaction with Tb927.7.3040 was lost (Fig 3B, green highlight). This raises the possibility that complex formation might depend also on Tb927.11.4900 GTP binding.

### Depletion of ZC3H5 complex components inhibits cytokinesis

If the three additional proteins form a complex with ZC3H5, they should all decrease expression when tethered. Previous screening results using random fragments suggested no such activity (2), so we checked the individual full-length proteins. Repression was indeed seen for two of three clones (Fig 3D, Supplementary Fig S6). Consistent with this, results of a preliminary experiment suggested that all four proteins migrate predominantly in the free fractions of a sucrose gradient, rather than with monosomes or polysomes (Supplementary Fig S7). However, the association of a minor fraction with monosomes or polysomes cannot be ruled out.

We next examined whether the three putative complex components were required for cell proliferation. In each case, we used cells in which one copy of the relevant gene had an in situ V5 tag at the N-terminus. RNAi against any of the three proteins prevented proliferation and caused accumulation of cells with 4N, and then >4N DNA content, indicating defective cytokinesis (Fig 4). In each case also, the culture recovered as cells without RNAi were selected. Loss of each protein therefore had an effect similar to that seen after ZC3H5 depletion.

**Fig 4.**
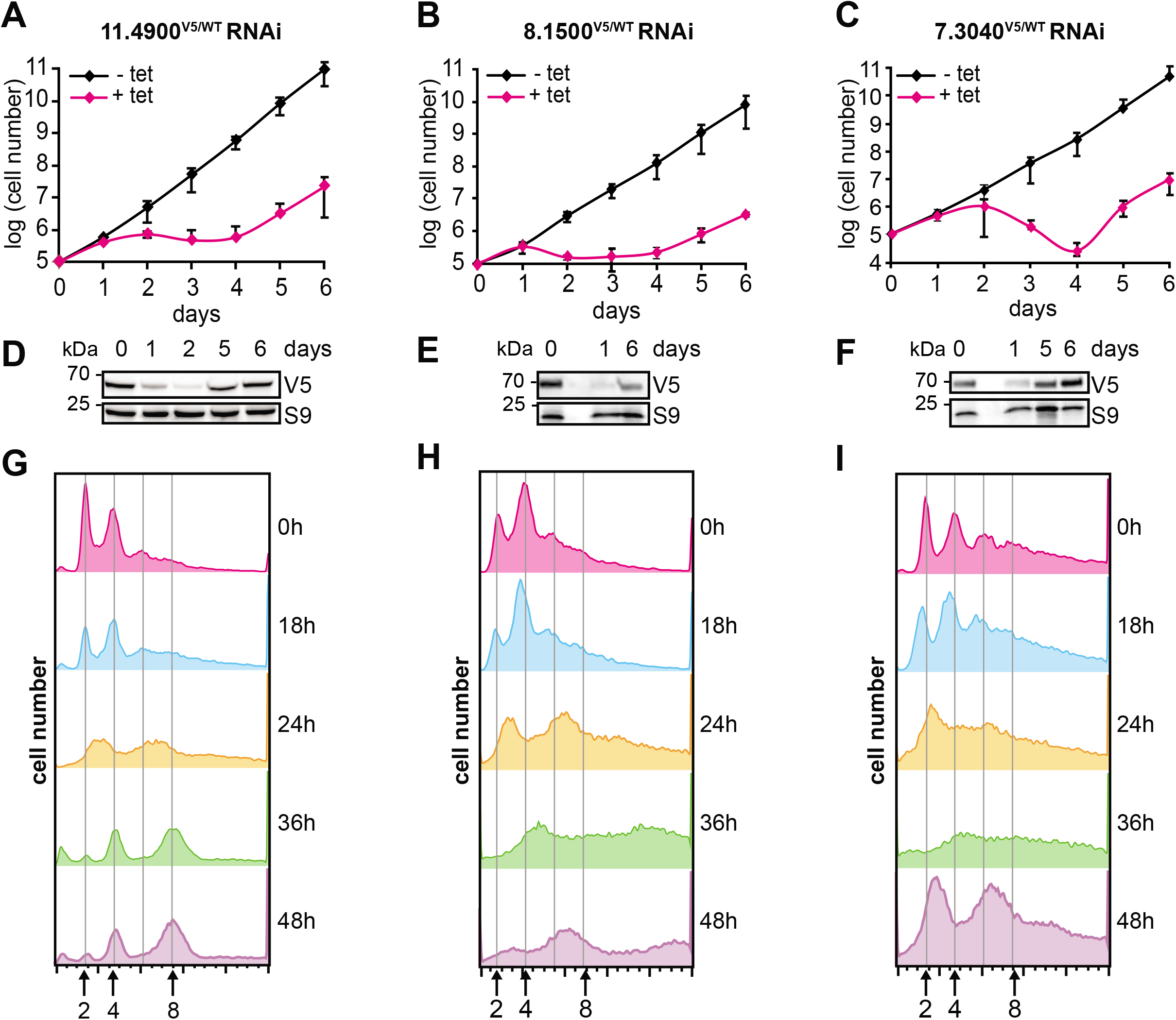
Effects of depleting ZC3H5 interaction partners. (A)-(C) Effects of depletion on cell growth. Cumulative cell numbers with and without tetracycline addition are shown for 3 replicates, with standard deviation. (D)-(F) Disappearance (and reappearance) of V5-tagged proteins after RNAi. In the cell line used, one copy was in situ V5-tagged and the other was intact. S9 served as a control. The number of days after tetracycline addition to induce RNAi are shown. (G)-(I) Effect of protein depletion on DNA content. DNA was stained with Propidium iodide and the cells were analyzed by FACS. The numbers below the diagrams indicate ploidy.

### ZC3H5 mRNA binding

To further examine the function of ZC3H5 we pulled down TAP-ZC3H5, released the RNA-protein complexes by TEV protease cleavage and sequenced RNA from both bound and unbound fractions. Principal component analysis (Supplementary Fig S8A) demonstrated clear separation of the bound and unbound fractions. The *ZC3H5* mRNA was more than 20-fold enriched in the bound fractions, consistent with selection via the nascent polypeptide (Supplementary Table S2, sheet 1). The trypanosome genome contains many repeated genes; if these are all counted, they can bias analysis. We therefore considered only a list of unique genes (13). Also, since mRNAs encoding ribosomal proteins are exceptionally short and behave abnormally in a variety of ways, they were omitted from the subsequent statistical analyses (Fig 5 and 7). There was a moderate correlation between binding and mRNA length (Fig 5A and Supplementary Fig S9). The 184 mRNAs that were reproducibly at least 4-fold enriched were significantly longer than mRNAs that were less than one-fold enriched (Fig 5A), suggesting a degree of non-specificity in the selection.

**Fig 5.**
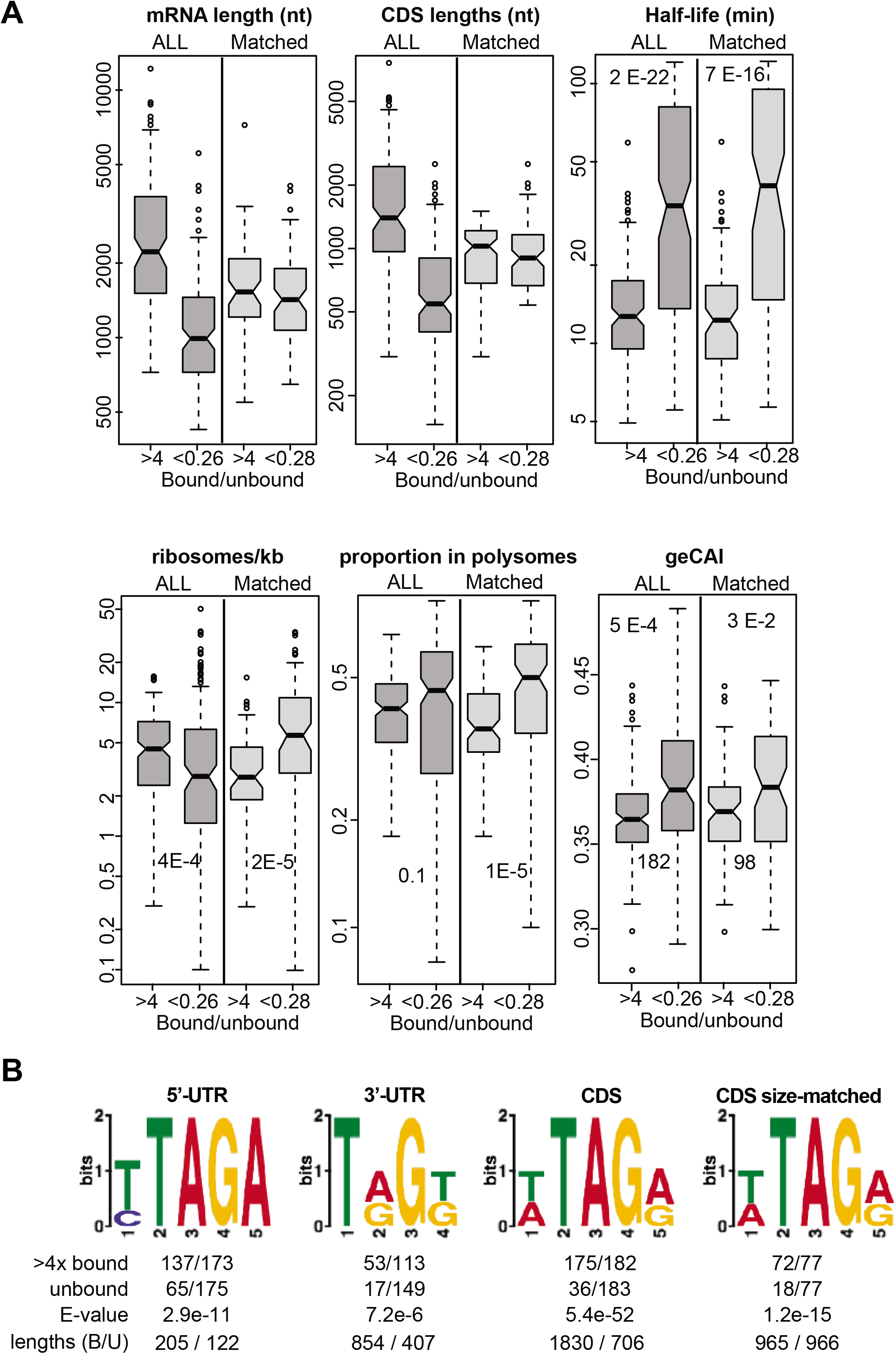
Binding of mRNAs by ZC3H5. (A) Comparison of bound and unbound mRNAs. The 182 “bound” mRNAs that showed at least 4-fold enrichment were compared with the 182 mRNAs that showed least binding, after exclusion of mRNAs encoding ribosomal proteins. These are labeled “ALL” in the plots. To eliminate possible bias caused by length differences, 98 mRNAs were selected from both sets in order to obtain matched distributions of coding region lengths; these results are labeled “Matched”. The genes concerned are listed in Supplementary Table S2, sheet 4. Half-lives are from (48), ribosome densities are from (49), the proportion in polysomes is from this paper, and the geCAI values are from (32). Values for all genes are shown as scatter plots in Supplementary Fig. S7. (B) Motifs enriched in the bound mRNAs as compared with the unbound mRNAs as in (A). 5’-UTRs, 3’-UTRs and coding sequences (CDS) were considered separately; some UTRs are not annotated. “>4xbound” and “unbound” show the number of sequences containing the motif relative to the total number considered. The E-value is from DREME. The median lengths of the sequences considered are shown at the bottom, with bound first and unbound second. Results for the matched CDSs are shown on the far right.

To find motifs that are enriched in bound mRNAs we compared the 184 mRNAs that were at least 4-fold enriched with a similar number of mRNAs that were least enriched (again excluding ribosomal protein mRNAs) (Supplementary Table S2, sheet 4). The 5’-UTRs, 3’-UTRs and coding regions were compared separately; most of the 5’-UTRs are annotated but only 60% of mRNAs have annotated 3’-UTRs. For all three comparisons, there was significant enrichment of motifs including the sequence UAG (Fig 5B). However, this might have been an artifact because the average sequence lengths examined from the bound fraction were 1.6-fold (UTRs) and 2.6-fold (coding regions) longer than those from the unbound fraction (Fig 5A). We therefore compared a manually-selected subset of bound and unbound coding regions that were as closely length-matched as possible (Supplementary Table S2, sheet 4). Enrichment of (U/A)UAG(A/G) persisted (Fig 5B), suggesting that ZC3H5 indeed preferentially selects mRNAs containing the motif.

We next looked at the biological characteristics of the bound mRNAs. There was no correlation between binding and mRNA half-life (Supplementary Fig S9B), but it was nevertheless notable that there were no mRNAs with half-lives above 120 min in the ZC3H5-bound fraction. Comparison of the bound and unbound sets also revealed a statistically significant preference for association with mRNAs with shorter half-lives (Fig 5A). The correlation between binding and ribosome occupancy was probably too weak to be meaningful (Supplementary Fig S9C, D and Fig 5A). There were no significant differences in the numbers of upstream open reading frames, or in open reading frame codon optimality (Fig 5A), between bound and unbound RNAs. When we examined functional classes, however, there was clear preferential binding of mRNAs from genes that are related to PAG genes (Fig 6A). PAG genes (procyclin-associated genes) were originally found downstream of, and co-transcribed with, the genes encoding the major procyclic surface proteins, the procyclins (14). There are however various other PAG orthologues that are transcribed by RNA polymerase II, are easily distinguished from the procyclin-associated genes at the DNA sequence level, and are well expressed in bloodstream forms. Seven PAG genes are included in the set of unique genes and mRNAs from all of them were enriched in the ZC3H5-bound fraction. Scrutiny of transcriptome data, as well as some Northern blot data for one of the relevant genes (15), suggested that these mRNAs are mostly less than 2 kb long, so they were not selected because they are inordinately long. Each of the open reading frames has at least 3 copies of the potential (U/A)UAG(A/G) motif.

As an independent verification of the motif we looked for it in the open reading frames encoding ribosomal proteins, since these had been excluded from the motif search. These sequences comprise, in total, over 32kb; from their base composition, and allowing for the absence of in-frame UAG stop codons in open reading frames, over 70 (U/A)UAG(A/G) motifs would be expected. In contrast, only two of the coding regions contain the motif, and one of those mRNAs was not depleted in the bound fraction.

### Effects of ZC3H5 depletion on the transcriptome and translation

Since reporter expression is inhibited by tethered ZC3H5, we expect its depletion to increase the abundance and/or translation of bound mRNAs. We tested this by characterizing the transcriptomes of parasites after 9h and 12h RNAi (Supplementary Table S2). These two time-points were chosen because the cytokinesis defect was just beginning to be apparent by 12h and we hoped to avoid secondary effects. We compared the results with those from wild-type cells because the uninduced cells already had a slightly unusual cell cycle profile (Fig 1). Principal component analysis showed good separation of the samples (Supplementary Fig S8B) and ZC3H5 mRNA was significantly decreased at both time-points (Fig 7A, Supplementary Table S2). Moreover, results from 9h and 12h RNAi correlated well, with slightly increased effects at 12h (Fig 7A). Most of the significant changes were however extremely small (1.5-2-fold, Fig 7A-B). Although we confirmed four of them by Northern blotting, with increased effects at 18h (Fig 7C), interpretation of such small effects is difficult. Unexpectedly, ZC3H5-bound mRNAs were mostly slightly (but not significantly) decreased whereas, as expected, unbound mRNAs were, on average, unaffected (Fig 7D). The most marked and significant effects were increases in mRNAs encoding ribosomal proteins, which were the least bound class (Fig 6A, B and Fig 7B). Since these mRNAs are very stable, they are partially responsible for the slight positive correlation between the fold change and mRNA half-life (Fig 7E). We also looked at the proportions of each mRNA that are associated with polysomes in wild-type cells, using data from Supplementary Table S3 (see below). The mRNAs that increased were significantly more polysome-associated than those that decreased; which is in accordance with their longer halflives (translation is known to influence the mRNA half-life) Interestingly, there were significant but small increases (FDR=0.05, >1.5-fold) in mRNAs encoding five components of the Tric complex, five translation factors, six RNA-binding proteins and two components of the CAF1-NOT complex. Overall, the data suggested slight upregulation of mRNAs encoding the translation machinery (Fig 6B).

**Fig 6.**
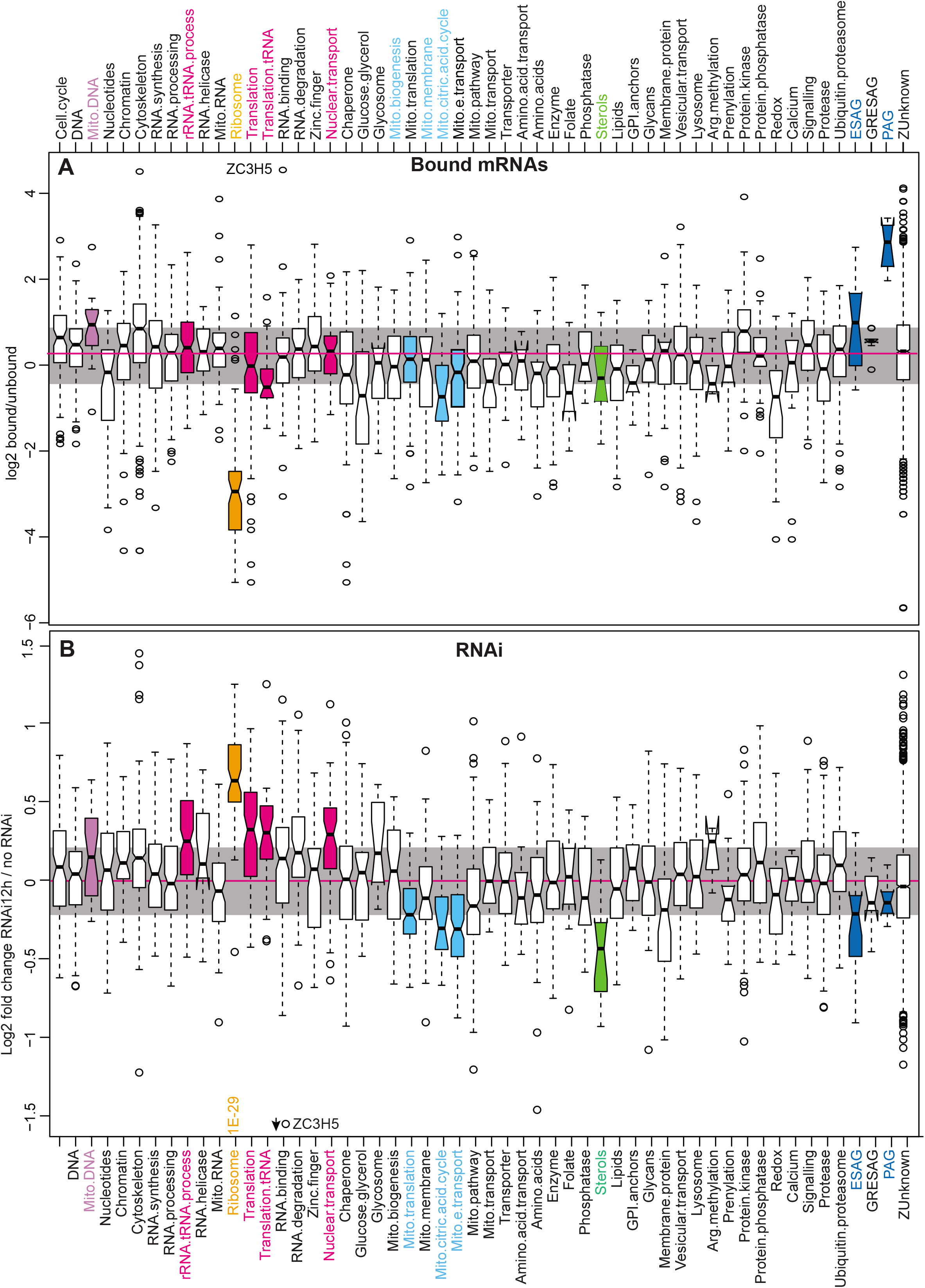
mRNAs associated with ZC3H5, and effects of ZC3H5 depletion. functional classes of encoded proteins. (A) The average of bound/unbound RNA for both replicates was calculated. All mRNAs were classified according to their protein function, where known, as in Supplementary Table S2. The ratios for each class are shown as box plots. Central line: median; upper and lower limits are 75th and 25th percentiles; Whiskers extend to the furthest measurement up to 1.5x the inter-quartile range; and circles are outliers. The shaded region shows the overall inter-quartile range and the overall median is in magenta. The ribosome class (significantly depleted by ANOVA, P=0.0009) is in brown and three classes that had medians above the overall 75th percentile are in cyan; only the PAG class is significantly enriched (P=0.02). (B) The fold changes on mRNA after 12h RNAi were calculated using DESeq2. All mRNAs were classified according to their protein function, where known, as in Supplementary Table S2. The ratios for each class are shown as box plots. Central line: median; upper and lower limits are 75th and 25th percentiles; Whiskers extend to the furthest measurement up to 1.5x the interquartile range; and circles are outliers. The shaded region shows the overall inter-quartile range and the overall median is in magenta. The ribosome class (significantly increased by ANOVA, P=1E-29) is in brown and other classes are colored if they have medians above the overall 75th percentile (pink) are or below the 25th percentile (mitochondrial in cyan, sterol metabolism in green). ANOVA P-values are indicated.

**Fig 7.**
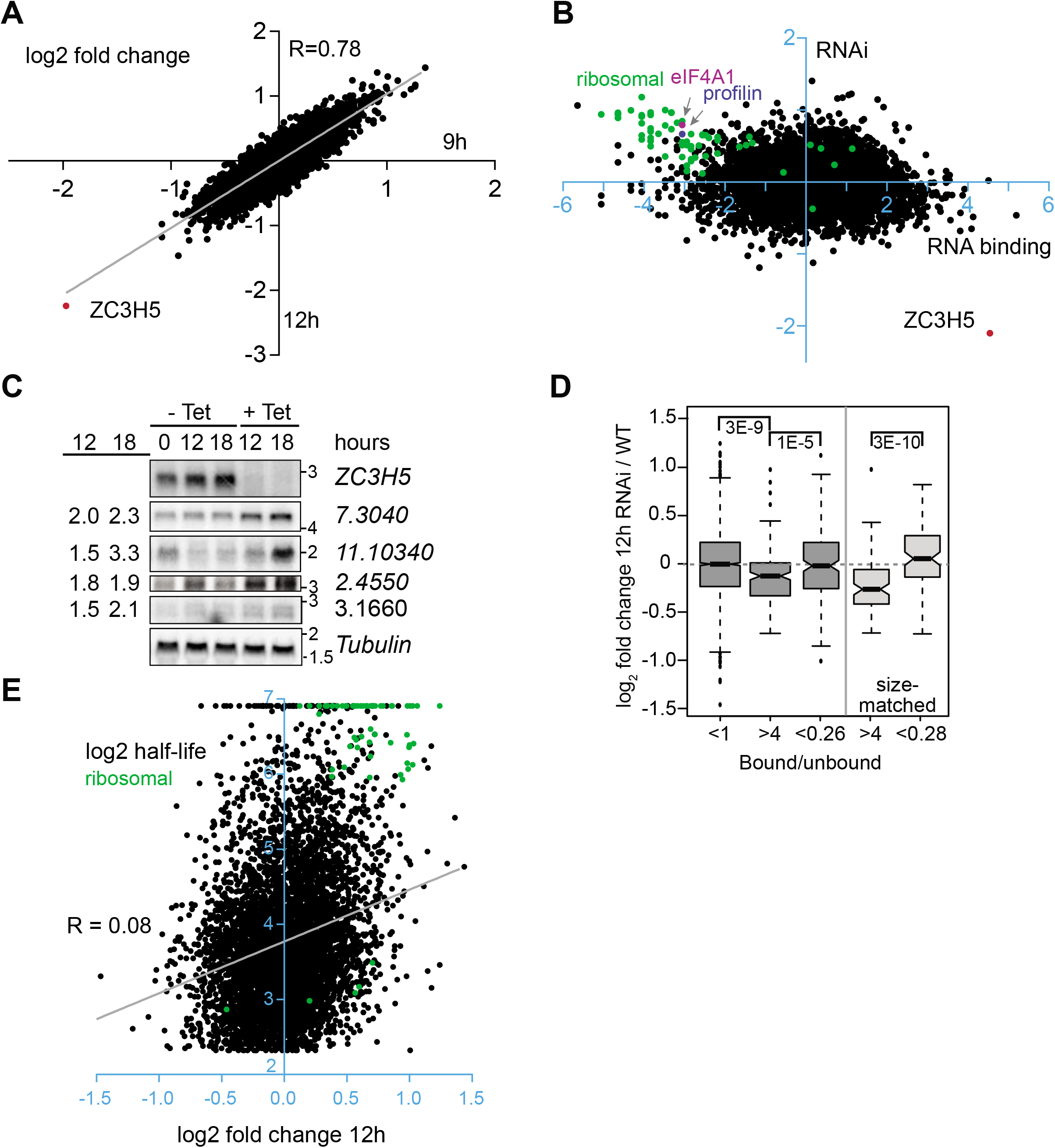
Effects of depleting ZC3H5 on the transcriptome. RNAi was induced in duplicate cultures of bloodstream-form trypanosomes for 9h or 12h, then RNA was analyzed by RNA-Seq. Results for a set of coding regions representing all unique genes are shown. Fold changes relative to duplicate uninduced cultures were calculated using DeSeq2 (45). (A) Log2 fold changes, relative to control, after 9h and 12h RNAi, calculated using DeSeq2. Each spot represents a single mRNA. (B) The average RNA binding ratio is on the x-axis and log2 fold changes, relative to control, after 12h RNAi on the y-axis. All values are log2 transformed. (C) Northern blot confirming increases in four mRNAs after ZC3H5 RNAi. Tubulin serves as a control. The numbers on the left indicate the signal intensity after 12 or 18h of RNAi, divided by the average signal intensity for cells grown without tetracycline, normalized to the tubulin control. (D) Effects of RNAi on mRNAs classified according to ZC3H5 binding. The box plot shows the median with 25th and 75th percentiles. Whiskers extend to the furthest point within 1.5-fold the inter-quartile range, and dots are outliers. “<1” are all mRNAs with a maximal binding ratio of less than 1 in both replicates; “>4” are the 183 mRNAs with a minimal binding ratio of 4 in both replicates. “<0.26” are the 183 mRNAs that showed lowest binding, excluding mRNAs that encode ribosomal proteins. The size-matched datasets (lighter grey fill) are 98 lowest-binding mRNAs selected such that their coding region lengths match those of 98 mRNAs that showed >4x binding (see supplementary Table S2). Results of pair-wise t-tests are shown above the boxes. (E) Half-lives of mRNAs relative to the fold change at 12h. Ribosomal protein mRNAs are in green.

Since the results from total mRNA were somewhat paradoxical, we wondered whether translation was selectively affected. We therefore examined the effect of *ZC3H5* RNAi on the polysomal distribution of mRNAs, subjecting extracts to sucrose gradient centrifugation. The polysome profiles were altered, with an increase in monosomes at the expense of the heaviest polysome fractions (Fig 8A, B; another example in Supplementary Fig S10A-C, upper panels). A similar change is seen in stumpy-form trypanosomes (16) but the transcriptome changes upon ZC3H5 depletion showed no correlation with those seen during stumpy-form differentiation (Supplementary Fig S10D).

**Fig 8.**
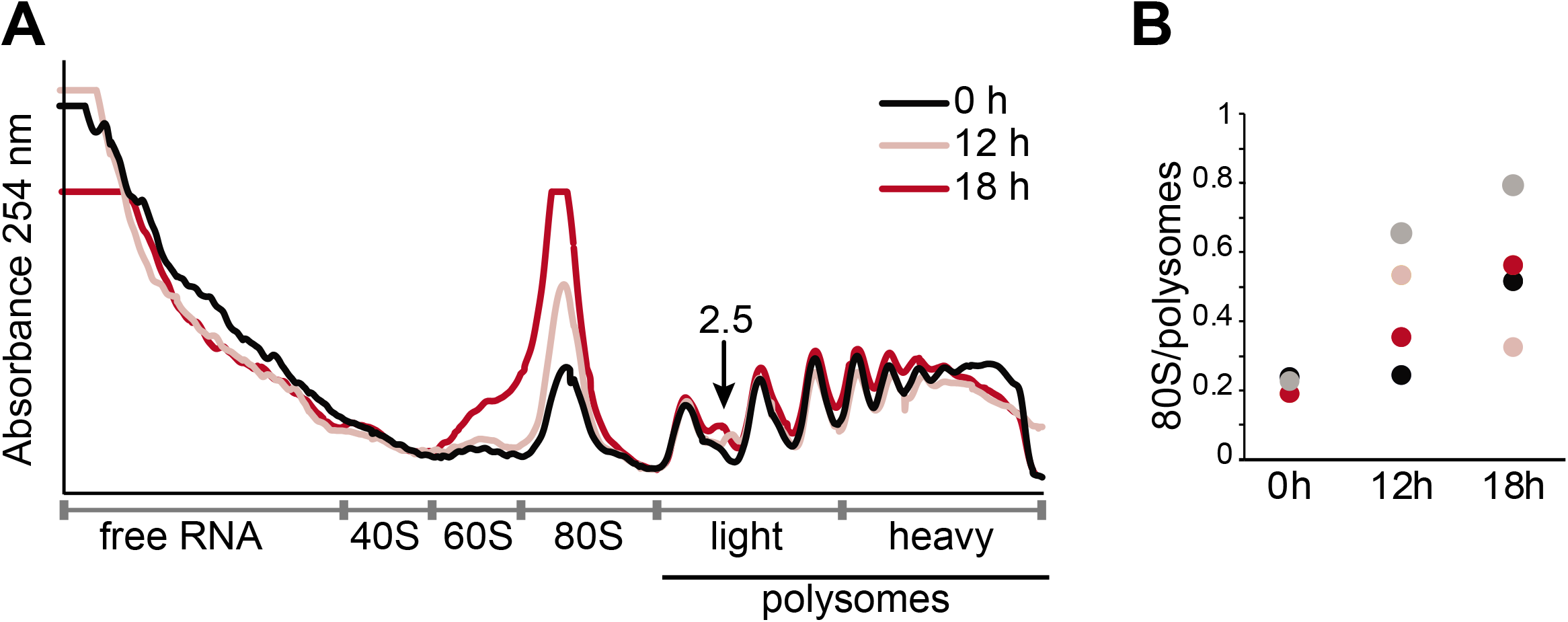
Effects of depleting ZC3H5 on polysomes. Ribosomes and polysomes were separated on sucrose gradients, after 12h, 18h RNAi or without RNAi (0h). (A) The absorbance at 254nm (random units) is shown for one experiment. The arrow points to the additional “2.5x” peak after RNAi; it suggests a defect in commencing elongation. (B) Quantitation of the ratio between the 80S and polysome 254 nm measurements for four independent experiments.

To find out whether specific mRNAs had moved into the monosome fraction after ZC3H5 depletion, we sequenced the mRNAs in free, monosome, light polysome (2,3 and 4 ribosomes) and heavy polysome (>4 ribosomes) fractions. This analysis can show major movements of mRNAs between nontranslated and translated fractions, but is too insensitive to detect more subtle alterations. Movement of *ZC3H5* mRNA from the polysome fractions to the free fraction was indeed clearly demonstrated (Supplementary Fig S8C). However, the principal component analysis suggested no significant changes in mRNA distributions between the fractions after RNAi (Supplementary Fig S8D). Some statistically significant differences for some mRNAs in the free fractions (Supplementary Table S3), can be ignored because they were for mRNAs that were predominantly in the other fractions. Otherwise, no significant differences were seen. These results suggest that the increase in monosomes probably represented free ribosomes. Despite this, there was no effect on the total mRNA amount (Supplementary Fig S11A) or on the overall protein synthesis pattern, at least for the more strongly-translated mRNAs (Supplementary Fig S11B).

Interestingly, the polysome profiles of ZC3H5-depleted trypanosomes had a clear new peak at the “2.5” position. “Halfmers” in polysome profiles consist of 1 or more full ribosomes plus a 40S subunit (17), and they arise when there are problems with 60S subunit joining. One possible cause is defective large subunit biogenesis (18). In trypanosomes, defects in ribosome biogenesis and export from the nucleus can cause feed-back inhibition of rRNA processing (19-21). A role of the ZC3H5 complex in ribosome biogenesis seemed unlikely, given its cytoplasmic location, and indeed, ZC3H5 depletion had no reproducible effect on the overall pattern of rRNAs and precursors (Supplementary Fig S12).

## Discussion

ZC3H5 co-purifies with three other proteins and our results suggest that they act as a complex. Each one inhibits expression when tethered in the 3’-untranslated region of a reporter mRNA, and depletion of any one of the four subunits impairs cytokinesis.

Tb927.11.4900 protein has a circularly permuted GTPase domain. Previously characterized eukaryotic (non-organellar) proteins with this domain arrangement - typified by yeast Noa1, Lsg1, Gnl1, Gnl2 and Gnl3l (22) - are mostly in the nucleolus and are implicated in ribosome biogenesis (23). These proteins are conserved in eukaryotic evolution and homologues are indeed present in trypanosomes. Because of its location, Tb927.11.4900 is unlikely to have a role in nuclear ribosome biogenesis, but it (or the complete complex) might be implicated in cytoplasmic ribosome maturation or modification. Mutation of the Tb927.11.4900 GTP binding domain prevented interaction with the Tb927.7.3040 protein in the two-hybrid assay, suggesting that complex formation might be regulated by GTP-GDP exchange.

The weak interactions observed with the potential Sdo1 homolog and with a possible GTPase-activating protein (Tb927.7.2470), open up the possibility that these two proteins regulate formation of the ZC3H5 complex. Yeast Sdo1 is a GTP-GDP exchange factor that is involved in the final stages of ribosome quality control: it releases Tif6 (eIF6 in human) from 60S ribosomes in the cytoplasm, allowing them to associate with 40S subunits (24) by activating the classical GTPase Ria1 (25) *(T. brucei* Tb927.3.2170). Sdo1 is conserved as far as archaea; the human equivalent is the protein implicated in Shwachman-Bodian-Diamond Syndrome (24). Thus, if these two proteins play a role on formation of the ZC3H5 complex, they may also link the complex to late ribosome maturation.

Loss of ZC3H5 caused a slight decrease in very heavy polysomes, and an increase in monosomes. It is normally assumed that 80S ribosomes dissociate into their component subunits upon translation termination. If this were universally true, the 80S peak would represent mRNAs bearing a single ribosome each. Indeed, in growing *S. cerevisiae*, most 80S ribosomes are elongating (26). Free monosomes have however been observed in mammalian cells subject to ribotoxic stress (27). An increase in monosomes - but not 2.5mers - is seen in stumpy-form trypanosomes (16). Since these also show a strong decrease in mRNA levels, it seems likely that free 80S subunits are present. We observed no movement of mRNAs towards the monosome fraction in ZC3H5-depleted cells, so we suggest that their monosomes are not associated with mRNAs. It is impossible to tell whether this is a direct effect of ZC3H5 depletion, or a secondary symptom of the onset of growth inhibition.

The appearance of 2.5mers in ZC3H5-depleted cells is a much more specific defect, and has not been observed previously in trypanosomes. In budding yeast, halfmers were first seen after treatment with low concentrations of cycloheximide (17), with clear peaks at the 1.5, 2.5, 3.5 and 4.5mer positions. Halfmers were also seen after depletion of proteins required for ribosome biogenesis, export, cytoplasmic maturation or translation initiation (18,28-31); they indicate a failure of 60S ribosomal subunit joining. In ZC3H5-depleted cells, however, the only increase detected was in 2.5mers, arguing against a general late ribosome biogenesis defect. The 2.5mers could consist of one ribosome on the start codon, a second 80S ribosome immediately downstream of the first and a 40S subunit “queueing” behind them in the 5’-UTR. This could be caused by a newly initiated ribosome failing to continue elongation. The coding regions and 5’-UTRs of mRNAs that bound to ZC3H5 were enriched in the motif [U/A]UAG[U/A]. Since UAG is a stop codon, this suggests an increased presence of four codons that specify amino acids: [U/A]UA (perhaps followed by G) and AG[U/A] (perhaps preceded by U). Comparison of the first 60 coding nucleotides of the 182 bound mRNAs (4x) with the same number of least enriched mRNAs still showed significant (1.1E-7) enrichment of UAG, present in 80 of the bound but only 22 of the unbound mRNAs. Using the codon optimality indices calculated in (32), the codons xUA and AGx are, on average, sub-optimal (Supplementary Fig. S13).

After ZC3H5 depletion, the vast majority of bound mRNAs showed no change in abundance, and a few decreased. This is the reverse of the effect that would be expected if the ZC3H5 complex suppresses expression. We therefore suspect that the tethering results are misleading. This could be because the complex does not work in the normal way when attached to the 3’-UTR, or because its effects are dependent on the nature of the open reading frame. We know that slow translation elongation can favor correct folding of proteins that are prone to aggregation (33), and therefore more active product than when translation is faster However, so far as we know, the failure of a protein to fold does not normally result in a decrease in the amount of the encoding mRNA. We propose, instead, that the codon composition of the reporter influences the effect of ZC3H5 complex binding. This is suggested by the fact that motifs associated with ZC3H5 binding are enriched in the coding region. In yeast, increases in the initiation rate have different effects on translation depending on the codon optimality of the open reading frame (34). If the coding region is easily translated, increasing the initiation rate will increase the translation rate. In contrast, if there are clusters of sub-optimal codons, increasing the initiation rate will lead to ribosome collisions, which result in premature termination and RNA cleavage by the quality control machinery (34). Thus, a complex that decreases translation of an optimal open reading frame might actually enhance translation of an open reading frame with sub-optimal codons.

In summary, we propose that the ZC3H5 complex is implicated in quality control during the translation of sub-optimal open reading frames, and that its assembly may be regulated by a GTP-GDP hydrolysis and exchange cycle.

## Experimental procedures

### Cells and plasmids

All experiments were done with Lister 427 monomorphic bloodstream-form parasites expressing the Tet-repressor. Parasites were cultivated in HMI-9 medium in an incubator at 37°C with 5% CO2. Stable cell lines were made with in situ TAP-, V5- or YFP-tagged ORFs. Most of the plasmids were assembled using the NEBuilder HiFi DNA assembly mastermix (NEB) following manufacturer’s recommendations. RNAi target gene fragments were selected based on default settings of the RNAit software (35). They were amplified from genomic DNA using primers with a 5’-extension including *attB* sites for cloning into the stem-loop pGL2084 plasmid (36). The resulting plasmid was digested with BamHI and HindIII and the fragment containing the stem loop was subcloned into the pHD1146 plasmid. For the tethering assays, cell lines constitutively expressing CAT reporter with boxB actin 3’-UTR were cotransfected with plasmids encoding Tb927.8.1500, Tb927.7.3040 and Tb927.11.4900 in fusion with the lambda N-peptide (2,3). Stable transfectants were selected in the presence of appropriate selective drugs at the following concentrations: 1 μg ml-1 puromycin, 2.5 μg ml-1 phleomycin (InvivoGen), 5 μg ml-1 hygromycin B (Calbiochem) and 10 μg ml-1 blasticidin (InvivoGen) and independent clones obtained by limiting dilution. For yeast 2-hybrid, all entry clones were shuttled into the Gateway-compatible vector (37) and pair-wise analysis performed as previously described in (3). The Tb927.8.1500 and Tb927.11.4900 mutants were generated using the Q5 site-directed mutagenesis kit (NEB) from the wildtype vector templates. Details of all plasmids and oligonucleotides are found in Supplementary Table S4.

### Protein Analysis

Proteins were analyzed by Western blotting, and detected by enhanced chemiluminescence according to the manufacturer’s instructions (Amersham). Antibodies were against the V5 tag (AbD seroTec, 1:1000), the YFP tag (Roche, 1:1000) and aldolase (rabbit, 1:50000).

CAT was measured in a kinetic assay involving partition of 14C-buturyl chloramphenicol from the aqueous to the organic phase of scintillation fluid (38). Co-immunoprecipitations of endogenously V5-tagged ERBP1 (5), Tb927.8.1500, Tb927.7.3040 and Tb927.11.4900 proteins were done as previously described (39). Tandem affinity purification of *in situ* TAP-tagged ZC3H5 was essentially carried out according to (40). In each case, purified proteins were separated on SDS-4 to 20% polyacrylamide gradient gels (BioRad) and stained by colloidal Coomassie blue. Co-purified proteins and BF whole cell extracts were analyzed in three independent experiments by LC/MS by the ZMBH Mass Spectrometry facility. Raw data were analyzed using MaxQuant 1.5.8.3, with label-free quantification (LFQ) and match between runs enable (41). The identified proteins were filtered for known contaminants and reverse hits, as well as hits without unique peptides. Statistical analysis was performed in Perseus (42).

### RNA manipulation

Northern blots were probed with [32P]-labelled DNA from ZC3H5, Tb927.7.3040, Tb927.11.10340, Tb927.2.4550, Tb927.3.1660 and tubulin genes. To identify the transcripts bound to ZC3H5, 2×10^9^ cells were washed in ice-cold Voorheis’s-modified phosphate-buffered saline (vPBS; PBS supplemented with 10 mM glucose and 46 mM sucrose) and the cell pellet snap frozen in liquid nitrogen. The RNA immunoprecipitation was done essentially as described in (43). Briefly, the cell extracts were incubated with IgG-coupled magnetic beads (NEB) for 2 h at 4°C, and the unbound fraction was collected. After extensive washing, bound ZC3H5 was eluted using TEV protease for 2 h at 16°C. RNA was isolated using Trifast reagent (Peqlab). Total RNA from the unbound fraction was depleted of ribosomal RNA (rRNA) using RNAse H and a mixture of 50-base DNA oligos complementary to trypanosome rRNAs. The recovered RNA from both bound and unbound samples were then analyzed by RNA-Seq (David Ibberson, Bioquant). For transcriptome analysis, RNAi was induced with 250 ng.ml-1 tetracycline for either 9 or 12 h. Differences in mRNA abundance were assessed using DESeqU1 (44), a custom version of DESeq2 (45).

### Immunofluorescence microscopy and FACS analysis

Immunofluorescence of bloodstream cells was performed as previously described in (5). Primary antibodies were GFP 1:500 and tubulin 1:1000. Images were acquired using an Olympus IX81microscope and analyzed with Olympus xcellence software or image J. FACS analysis was performed as described in (46).

### Polysome fractionation and analysis

Polysome fractionation was performed according to (47). For each gradient, 5×10^8^ cells were collected by centrifugation at 3000 rpm for 15 min. The cell pellet was resuspended in 50 ml serum-free medium and transferred to a 50 ml conical tube. The cells were then incubated with 100μg/ml cycloheximide for 7 min at RT. Cells were pelleted again by centrifugation at 2300 rpm for 7 min at 4°C, washed with 1 ml ice-cold 1x PBS and transferred to an Eppendorf tube. Cells were then lysed in 350 μl lysis buffer (20 mM Tris-HCl pH 7.5, 20 mM KCl, 2 mM MgCl2, 2 mM DTT, 1000U RNasin ribonuclease inhibitor (Promega), 10 μg/ml leupeptin, 0.2% IGEPAL, 200 mM sucrose, 100 μg/ml cycloheximide) and passed 15 times through the 21-gauge needle using a 1-ml syringe and then 15 times through the 27-gauge needle. The lysate was cleared by centrifugation at 15000 g for 10 min at 4°C in a microfuge. Salt concentration was adjusted to 120 mM KCl and the lysate was loaded on the 4 ml continuous linear 15-50% sucrose gradient. The gradients were centrifuged at 40000 rpm for 2h at 4°C in a SW 60 Ti swinging-bucket rotor (Beckman Coulter). Afterwards, 16 fractions with a volume of 300 μl were collected by fractionation with the UV/VIS detector (Teledyne Isco). For RNA purification, 900 μl TriFast was added to each tube and RNA was purified following manufacturer protocol. For each fraction, the reads for each open reading frame per million reads (RPM) were calculated. In order to be able to estimate the real distributions of each mRNA across the gradient, a small portion of each sequencing sample was subjected to Northern blotting, using a spliced leader probe to detect mRNA, and a human beta-globin probe to check RNA yield after preparation and handling. The spliced leader signals were first corrected for RNA yield; then, we calculated the proportion of the total spliced leader signal that was in each fraction. The RPM values were multiplied by these values in order to obtain the representation in each fraction. To eliminate numbers after the decimal point, enabling analysis using in-house R scripts, all values were multiplied by 1000. Overall, the proportions (mean and standard deviation) were free: 35±4%; monosomes 24±3%; light polysomes 24±4%; and heavy polysomes 14±3%. Some error is introduced by slight variations in the precise position of the boundaries, especially between light and heavy polysomes: the percentage in polysomes was 39±3%.

### Data availability

The mass spectrometry data are available via ProteomeXchange with identifier PXD019073. ArrayExpress accession numbers for the RNASeq data are as follows: ZC3H5 RNAi: E-MTAB-9088; TAP-ZC3H5 pull-down: E-MTAB-9084; polysomes: E-MTAB-8069.

## Acknowledgements

We thank Nina Papavisiliou for generously hosting KB during the latter part of this project. We also acknowledge Claudia Helbig and Ute Leibfried for technical support, David Ibberson of the BioQuant sequencing facility for cDNA library construction, and the Mass spectrometry and Imaging facilities (Dr. Holger Lorenz) of the ZMBH.

## Funding

This work was supported by the Deutsche Forschungsgemeinshaft (Cl112/24 to C.C.) and core support from the State of Baden-Württemberg.

## Conflict of interest

The authors declare that they have no conflicts of interest with the contents of this article.

